# A dysregulated stromal remodelling programme characterises prior anti-TNF failure in ulcerative colitis

**DOI:** 10.64898/2026.08.11.744139

**Authors:** John P Thomas, Tyler Wooldridge, Domenico Cozzetto, Nathalie Lambie, Hiromi Kudo, Aamir Saifuddin, Lejla Gul, Dezso Modos, Robert Goldin, Nik Matthews, Tamas Korcsmaros, Nick Powell

## Abstract

Prior anti-tumour necrosis factor (TNF) failure is associated with reduced efficacy of subsequent advanced therapies in ulcerative colitis (UC), but the biological basis of this treatment-refractory state remains unclear. We integrated clinical outcomes and baseline colonic transcriptomic data from UC patients in the UNIFI phase III trial programme with regulatory and signalling network inference, connectivity mapping, and single-cell-resolution spatial transcriptomics. Colonic transcriptomic analyses identified coordinated enrichment of extracellular matrix organisation, collagen remodelling and integrin-associated programmes, increased stromal cell representation and elevated inferred MAPK/EGFR activity in UC patients with prior anti-TNF failure. Causal network inference prioritised MAPK3 as a candidate regulator of this state, while connectivity mapping identified MEK/EGFR inhibitors as candidate perturbagens. MEK inhibition suppressed stromal pathways and reduced inferred MAPK/EGFR activity *ex vivo*. Spatial profiling of active UC and non-IBD colonic tissues localised these programmes to UC-enriched stromal niches. Ligand-receptor inference further identified reciprocal stromal-myeloid communication within these niches. Collectively, these findings define a stromal remodelling programme associated with prior anti-TNF failure and nominate MAPK/EGFR signalling as a potentially tractable component of treatment-refractory UC.

## INTRODUCTION

Monoclonal antibodies targeting the pro-inflammatory cytokine tumour necrosis factor (TNF) were the first biologic therapies to transform the management of moderate-to-severe ulcerative colitis (UC). In the landmark ACT 1 and ACT 2 trials, infliximab was more effective than placebo for inducing and maintaining clinical response, clinical remission and mucosal healing in patients with moderately to severely active UC^1^. The therapeutic landscape has subsequently expanded to include agents targeting integrin-mediated lymphocyte trafficking, IL-12/23 and IL-23 signalling, Janus kinase signalling and sphingosine-1-phosphate receptors^2^. Despite this expanding therapeutic armamentarium, anti-TNF agents remain widely used as first-line advanced therapies^3^, and a substantial proportion of patients either fail to respond or subsequently lose response^4^.

Importantly, prior anti-TNF failure is associated with attenuated efficacy of subsequent advanced therapies. In a post hoc analysis of GEMINI 1, the placebo-adjusted benefit of vedolizumab for achieving clinical response at week 6 was numerically greater in anti-TNF-naïve patients than in those with prior anti-TNF failure (26.4% versus 18.1%)^5^. Similarly, in the True North trial, the placebo-adjusted benefit of ozanimod for achieving clinical remission at week 10 was greater in anti-TNF-naïve patients than in patients with prior anti-TNF failure (15.5% versus 5.4%)^6^. This attenuation of subsequent treatment efficacy which we term as drug sequencing depreciation, represents an important clinical challenge. As anti-TNF agents are frequently positioned early in the UC treatment algorithm^4^, many patients receive alternative advanced therapies only after anti-TNF failure. Consequently, the patients with the greatest need for an effective alternative treatment are often those least likely to respond to subsequent therapies. The observation that this pattern extends across agents with distinct mechanisms of action suggests that prior anti-TNF failure may identify a broader treatment-refractory phenotype rather than resistance confined to TNF blockade.

The biological basis of this treatment-refractory phenotype remains poorly understood. Anti-TNF failure is heterogeneous and may arise through primary non-response, secondary loss of response, immunogenicity or pharmacokinetic mechanisms^7^. Patients with prior failure may also have longer-standing or more severe disease, which could account for their poorer subsequent outcomes. Alternatively, their mucosal inflammation may be sustained by a distinct molecular and cellular state that is not adequately targeted by currently available therapies. These possibilities are not mutually exclusive. Distinguishing greater accumulated disease burden from a biologically distinct treatment-refractory mucosal state is therefore important for understanding drug sequencing depreciation and identifying alternative therapeutic strategies.

To address this question, we analysed baseline clinical characteristics, colonic biopsy transcriptomes and week 8 induction outcomes from patients with moderate-to-severe UC enrolled in the UNIFI phase III trial programme of ustekinumab, an antibody targeting the shared IL-12/23 p40 subunit^8^. We first characterised the clinical phenotype associated with prior anti-TNF failure in this cohort. Using an integrated *in silico* approach involving pathway enrichment, cell type deconvolution, gene regulatory network and upstream signalling network inference, we then found that the transcriptional landscape associated with anti-TNF failure in UC is characterised by dysregulated stromal remodelling and EGFR/MAPK signalling. Connectivity mapping was used to identify pharmacological perturbations predicted to reverse this molecular phenotype, with the prioritised MAPK pathway subsequently evaluated using transcriptional responses to MEK inhibition in *ex vivo* UC explant cultures. Finally, the cellular and spatial localisation of the implicated molecular pathways was resolved using single cell-resolution spatial transcriptomics and spatial ligand-receptor inference. Through this multimodal approach, we define a stromal remodelling programme associated with prior anti-TNF failure, implicate MAPK/EGFR signalling as a potentially tractable component of this treatment-refractory phenotype, and localise this programme to organised stromal-immune niches within inflamed UC mucosa.

## RESULTS

### UC patients with prior anti-TNF failure have greater disease burden compared to anti-TNF naïve patients

We evaluated publicly available data^9^ originating from the UNIFI trial which comprised baseline clinical information for 537 participants with moderate-to-severe UC (defined as a total score of 6-12 on the full Mayo score) and micro-array transcriptomic data from baseline colonic biopsies prior to the administration of ustekinumab or placebo, as well as various clinical outcome measures at 8 weeks post-induction therapy. 275 UC patients in this cohort had previously failed anti-TNF therapy, whilst 262 were anti-TNF naïve.

Patients with prior anti-TNF failure had significantly longer disease duration (7.4 years in anti-TNF failure group vs 5.19 years in anti-TNF naïve group, p = 8.83e-6; Mann-Whitney U test), higher CRP (5.39 mg/L in anti-TNF failure group vs 3.94 mg/L in anti-TNF naïve group, p = 0.034; Mann-Whitney U test), and greater histological severity as assessed by the total Geboes score (13 in anti-TNF failure group vs 12 in anti-TNF naïve group, p = 0.002; Mann-Whitney U test) (Table 1). These findings indicate that prior anti-TNF failure was associated with a greater disease burden.

**Table 1.**
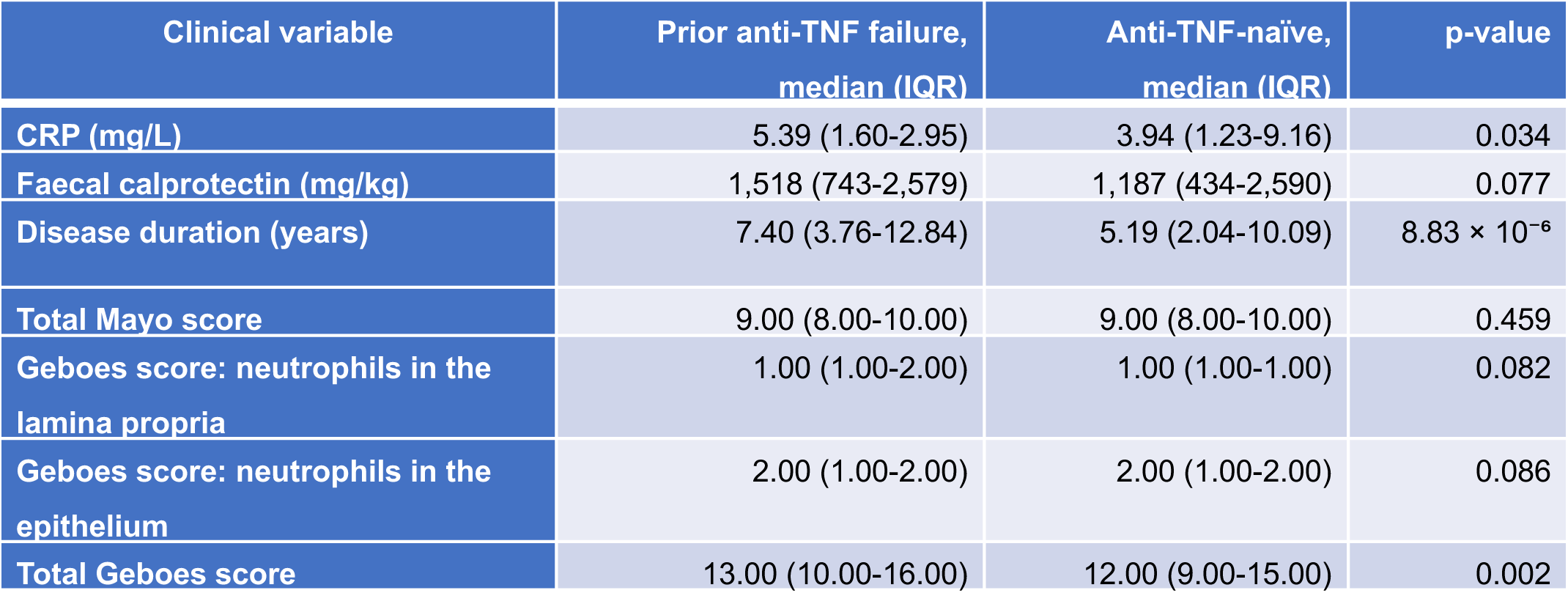

| Clinical variable | Prior anti-TNF failure,<br>median (IQR) | Anti-TNF-naïve,<br>median (IQR) | p-value |
| --- | --- | --- | --- |
| CRP (mg/L) | 5.39 (1.60-2.95) | 3.94 (1.23-9.16) | 0.034 |
| Faecal calprotectin (mg/kg) | 1,518 (743-2,579) | 1,187 (434-2,590) | 0.077 |
| Disease duration (years) | 7.40 (3.76-12.84) | 5.19 (2.04-10.09) | $8.83 \times 10^{-6}$ |
| Total Mayo score | 9.00 (8.00-10.00) | 9.00 (8.00-10.00) | 0.459 |
| Geboes score: neutrophils in the<br>lamina propria | 1.00 (1.00-2.00) | 1.00 (1.00-1.00) | 0.082 |
| Geboes score: neutrophils in the<br>epithelium | 2.00 (1.00-2.00) | 2.00 (1.00-2.00) | 0.086 |
| Total Geboes score | 13.00 (10.00-16.00) | 12.00 (9.00-15.00) | 0.002 |

### Prior anti-TNF failure is associated with worse week 8 induction outcomes to ustekinumab

We next examined whether prior anti-TNF failure was associated with worse induction outcomes specifically among patients receiving active ustekinumab treatment. Placebo-treated participants were excluded, and models were adjusted for baseline CRP, disease duration and total Geboes score, which differed between the anti-TNF-failure and anti-TNF-naïve groups. Of 357 ustekinumab-treated patients, 318 had complete data for the outcome and included covariates.

Prior anti-TNF failure was independently associated with lower odds of achieving clinical response at week 8 (adjusted OR 0.55, 95% CI 0.35-0.87; p = 0.010) and mucosal healing (adjusted OR 0.50, 95% CI 0.25-0.99; p = 0.047; Figure 1A; Table 2). Prior anti-TNF failure was also associated with lower odds of endoscopic healing (adjusted OR 0.56, 95% CI 0.31-1.02; p = 0.058) and clinical remission (adjusted OR 0.60, 95% CI 0.28-1.26; p = 0 .174) but these did not reach statistically significance.

**Figure 1:**
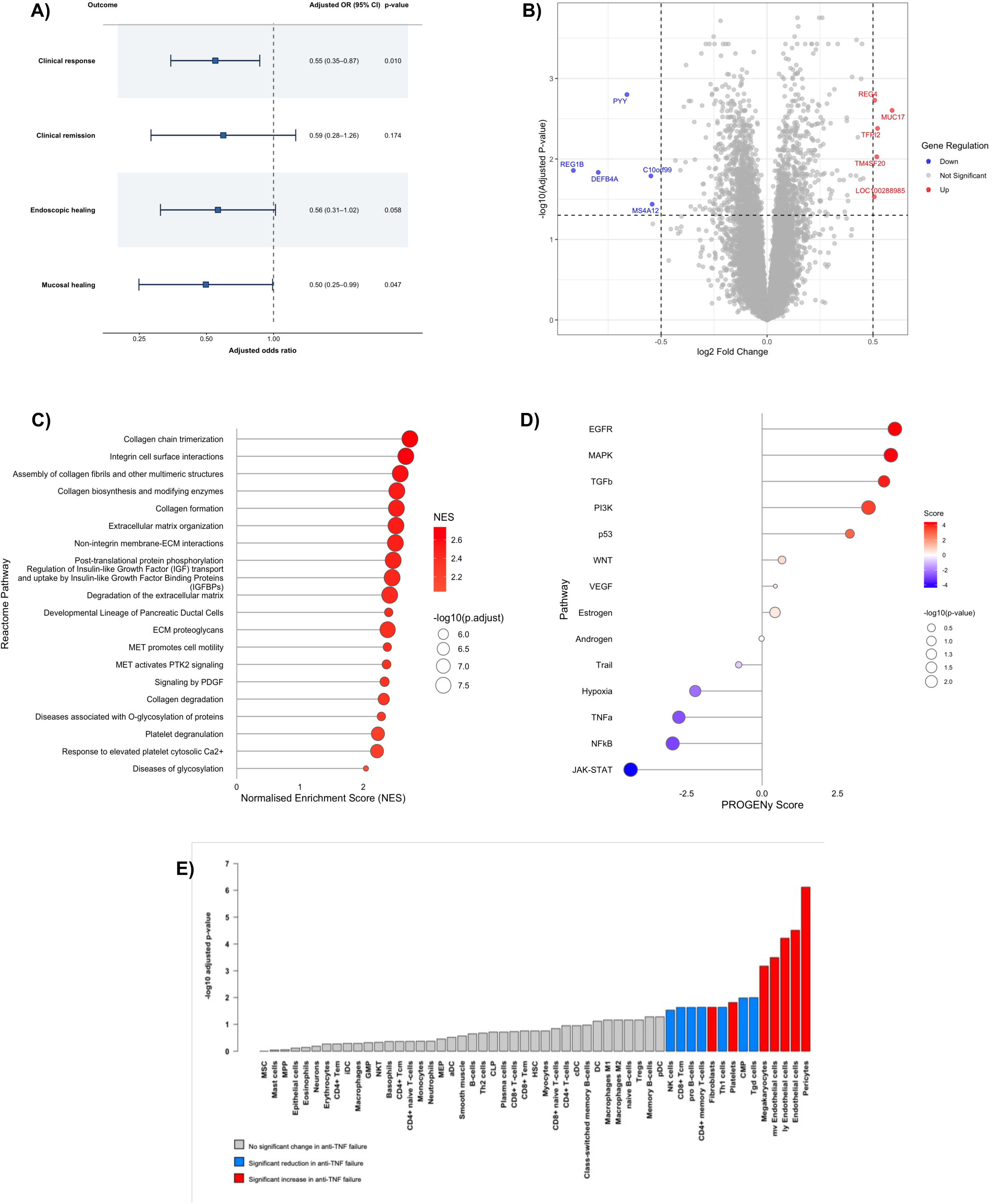
Clinical and molecular phenotype associated with prior anti-TNF failure in UC. **(A)** Forest plot showing the association between prior anti-TNF failure and week 8 induction outcomes among ustekinumab-treated participants. Separate multivariable logistic regression models were adjusted for log₂(CRP + 1), log₂(disease duration + 1) and total Geboes histological score. The analysis included 318 complete cases. Squares indicate adjusted odds ratios and horizontal lines indicate Wald 95% confidence intervals; the dashed vertical line denotes an odds ratio of 1. Anti-TNF-naïve participants were the reference group. **(B)** Volcano plot of adjusted differential gene expression in baseline colonic biopsies from patients with prior anti-TNF failure versus anti-TNF-naïve patients. Gene-wise linear models were adjusted for disease duration, Mayo Endoscopic Score, faecal calprotectin, CRP and total Geboes score. Positive log₂ fold changes indicate higher expression following prior anti-TNF failure. Genes meeting both the absolute log₂ fold-change threshold of 0.5 and Benjamini-Hochberg-adjusted *P* < 0.05 are shown in red (higher expression) or blue (lower expression); selected genes are labelled. **(C)** The 20 Reactome pathways with the highest positive normalised enrichment scores in patients with prior anti-TNF failure. Point size represents −log₁₀(Benjamini-Hochberg-adjusted *P* value), and colour represents the normalised enrichment score. **(D)** PROGENy-inferred pathway activity between UC patients with prior anti-TNF failure and anti-TNF-naïve patients. Positive scores indicate higher inferred activity following prior anti-TNF failure and negative scores indicate higher activity in anti-TNF-naïve patients. Point colour represents the normalised pathway-activity score and point size represents −log₁₀(*P* value); pathway *P* values were adjusted across pathways using the Benjamini-Hochberg method. **(E)** Comparison of xCell-derived cell-type enrichment scores between prior anti-TNF-failure and anti-TNF-naïve groups. Bars are ordered by statistical significance and their height represents −log₁₀(Benjamini-Hochberg-adjusted *P* value). Red denotes significantly higher and blue significantly lower inferred enrichment following prior anti-TNF failure; grey denotes no significant difference at false-discovery rate <0.05. Direction was determined from the difference in median xCell scores between groups. CRP, C-reactive protein; ECM, extracellular matrix; FDR, false-discovery rate; NES, normalised enrichment score; OR, odds ratio; UC, ulcerative colitis.

**Table 2.**

| Week 8 outcome | Patients analysed, N | Achieving outcome, N | Adjusted OR | 95% CI | P value |
| --- | --- | --- | --- | --- | --- |
| Clinical response | 318 | 165 | 0.55 | 0.35-0.87 | 0.010 |
| Clinical remission | 318 | 37 | 0.60 | 0.28-1.26 | 0.174 |
| Endoscopic healing | 318 | 65 | 0.56 | 0.31-1.02 | 0.058 |
| Mucosal healing | 318 | 46 | 0.50 | 0.25-0.99 | 0.047 |

Collectively, these findings indicate that prior anti-TNF failure identifies a subgroup of UC patients with a reduced likelihood of achieving favourable outcomes following ustekinumab induction. In particular, the associations with clinical response and mucosal healing persisted after adjustment for observed differences in disease duration, CRP and baseline histological severity.

### There are subtle differences in gene expression in the colonic mucosa of UC patients who failed anti-TNF therapy compared to anti-TNF naïve UC patients

To investigate the molecular phenotype associated with anti-TNF failure in UC patients, differential gene expression analysis was performed on microarray data derived from the baseline colonic biopsies of UC patients who previously failed anti-TNF therapy versus those who were anti-TNF naïve in the UNIFI cohort. The differential expression model adjusted for differences in baseline clinical characteristics (CRP, disease duration, total Geboes score, Mayo Endoscopic Score, and faecal calprotectin) between the two groups, enabling the identification of transcriptional differences associated with prior anti-TNF failure independently of these measured clinical characteristics.

This analysis revealed that 10 genes achieved a pre-specified threshold for differential expression of absolute log2 fold change >0.5 and adjusted p-value <0.05 (Figure 1B). These genes included those related to epithelial barrier function (*MUC17, DEFB4A*), epithelial regeneration and repair (*REG4, REG1B*), cell signalling and immune response (*TM4SF20, MS4A12*), appetite regulation (*PYY*), coagulation (*TFPI2*) as well as genes with limited functional characterisation (*C10orf99, LOC100288985*).

### Extracellular matrix and stromal pathways are highly enriched in the colonic mucosa of UC patients who failed anti-TNF therapy compared to anti-TNF naïve UC patients

Gene set enrichment analysis (GSEA) was then performed using the Reactome database to evaluate whether there may be differences in the enrichment of biological pathways between anti-TNF and anti-TNF naïve UC patients. This revealed multiple enriched Reactome pathways in anti-TNF failure (adjusted p value < 0.05) of which the top 20 according to the normalised enrichment score (NES) is depicted in Figure 1C. Pathways relating to the extracellular matrix (ECM) and stroma were highly enriched in UC patients who previously failed anti-TNF therapy compared to those who were anti-TNF naïve. These included Reactome terms such as extracellular matrix organisation, non-integrin membrane ECM interactions, integrin cell surface interactions, degradation of the extracellular matrix, L1CAM interactions, laminin interactions, ECM proteoglycans, and collagen formation.

### Prior anti-TNF failure is associated with a shift towards EGFR-MAPK signalling in the colonic mucosa

To investigate whether prior anti-TNF failure is associated with distinct mucosal signalling programmes, pathway activity was inferred from colonic transcriptomic data using PROGENy, a perturbation-based framework for estimating signalling pathway activity from gene expression profiles^10^.

Comparison of patients with prior anti-TNF failure versus those who were anti-TNF naïve revealed substantial differences in inferred mucosal pathway activity (Figure 1D). Prior anti-TNF failure was associated with lower inferred activity of JAK-STAT signalling (normalised PROGENy pathway activity score = −4.36, adjusted p = 0.004), NF-κB signalling (score = −2.96, adjusted p = 0.006), TNFα signalling (score = −2.76, adjusted p = 0.012) and Hypoxia-associated signalling (score = −2.22, adjusted p = 0.037). Conversely, higher inferred activity was observed for EGFR (score = 4.40, adjusted p = 0.004), MAPK (score = 4.27, adjusted p = 0.004), TGF-β (score = 4.04, adjusted p = 0.032) and PI3K signalling (score = 3.53, BH-adjusted p = 0.004).

Collectively, these findings indicate that prior anti-TNF failure is associated with a marked shift in the mucosal signalling landscape. Whereas anti-TNF-naïve patients exhibited greater activity of canonical inflammatory pathways centred on TNFα, NF-κB, and JAK-STAT signalling, anti-TNF failed patients demonstrated relative enrichment of growth factor receptor-associated pathways, including EGR, MAPK, and PI3K signalling. These observations suggest that anti-TNF failure may be associated with a distinct molecular state that extends beyond differences in overall inflammatory burden.

### Anti-TNF failure in UC is associated with stromal remodelling of the colonic mucosa

To identify the cellular populations underlying the transcriptional differences associated with prior anti-TNF failure, cellular deconvolution analysis was performed using xCell, which infers relative cell type enrichment from bulk transcriptomic profiles based on reference expression signatures^11^.

Application of this approach to the UNIFI cohort revealed that, overall, UC patients with prior anti-TNF failure exhibited a significant increase in stromal cell abundance compared with anti-TNF-naïve patients (StromaScore adjusted p = 1.03 × 10⁻⁴), whereas no significant difference was observed in overall immune cell populations (ImmuneScore adjusted p = 0.430) (Table 3).

**Table 3.**

|  | Anti-TNF naïve<br>(median score) | Anti-TNF failed<br>(median score) | Adjusted p-value |
| --- | --- | --- | --- |
| ImmuneScore | 0.689 | 0.659 | 0.430 |
| StromaScore | 0.052 | 0.069 | 1.03e-04 |

At the level of specific cell types, pericytes (adjusted p = 7.62 × 10⁻⁷), endothelial cells (adjusted p = 3.08 × 10⁻⁵), and fibroblasts (adjusted p = 0.023) were predicted to be significantly enriched in patients with prior anti-TNF failure compared to those who were anti-TNF-naïve (Figure 1D). These findings suggest that anti-TNF failure is associated with a shift in the stromal tissue compartment characterised by expansion of structural and vascular-associated stromal populations. In contrast, several T-cell populations were predicted to be reduced in anti-TNF-exposed patients, including CD4⁺ Th1 cells, CD4⁺ Th2 cells, naïve CD8⁺ T cells, and memory CD8⁺ T cells, suggesting relative depletion of adaptive immune cell populations within the mucosa.

To further evaluate cellular changes using an independent deconvolution framework, CIBERSORT which specifically focuses on immune cell populations was applied to the same transcriptomic dataset ^12^. Aligning with xCell results, CIBERSORT also demonstrated a significant reduction in CD8⁺ T cells (adjusted p = 0.0457) and γδ T cells (adjusted p = 0.0457) in patients with prior anti-TNF failure compared with anti-TNF-naïve individuals (Table 4).

**Table 4.**

|  | Anti-TNF failure group<br>(Median score) | Anti-TNF naïve<br>(Median score) | Adjusted p-value |
| --- | --- | --- | --- |
| CD8+ T cells | 0.059 | 0.070 | 0.0457 |
| Gamma delta T cells | 0.037 | 0.048 | 0.0457 |

Collectively, these analyses indicate that prior anti-TNF failure is associated with cellular remodelling of the colonic mucosa, characterised by an expansion of stromal compartments including pericytes, endothelial cells, and fibroblasts and a relative reduction in specific T-cell populations.

### Prior anti-TNF failure in UC is underpinned by transcriptional dysregulation

To investigate whether the observed cellular remodelling and pathway-level alterations associated with prior anti-TNF failure may be driven by changes in transcriptional regulation, transcription factor (TF) activity was inferred using VIPER^13^. This approach estimates TF activity based on the expression of downstream target genes defined in curated regulons. TF-target interactions were obtained from the DoRothEA database^14^.

VIPER analysis identified widespread alterations in transcriptional regulatory activity in UC patients with prior anti-TNF failure compared with anti-TNF-naïve individuals. In total, 84 transcription factors exhibited significant changes in activity (adjusted p < 0.05), comprising 71 with increased activity and 13 with decreased activity. Of these, 56 TFs demonstrated a high magnitude of effect (absolute normalised enrichment score > 3). The top 20 TFs by absolute normalised enrichment score as predicted by VIPER are depicted in Figure 2A.

**Figure 2:**
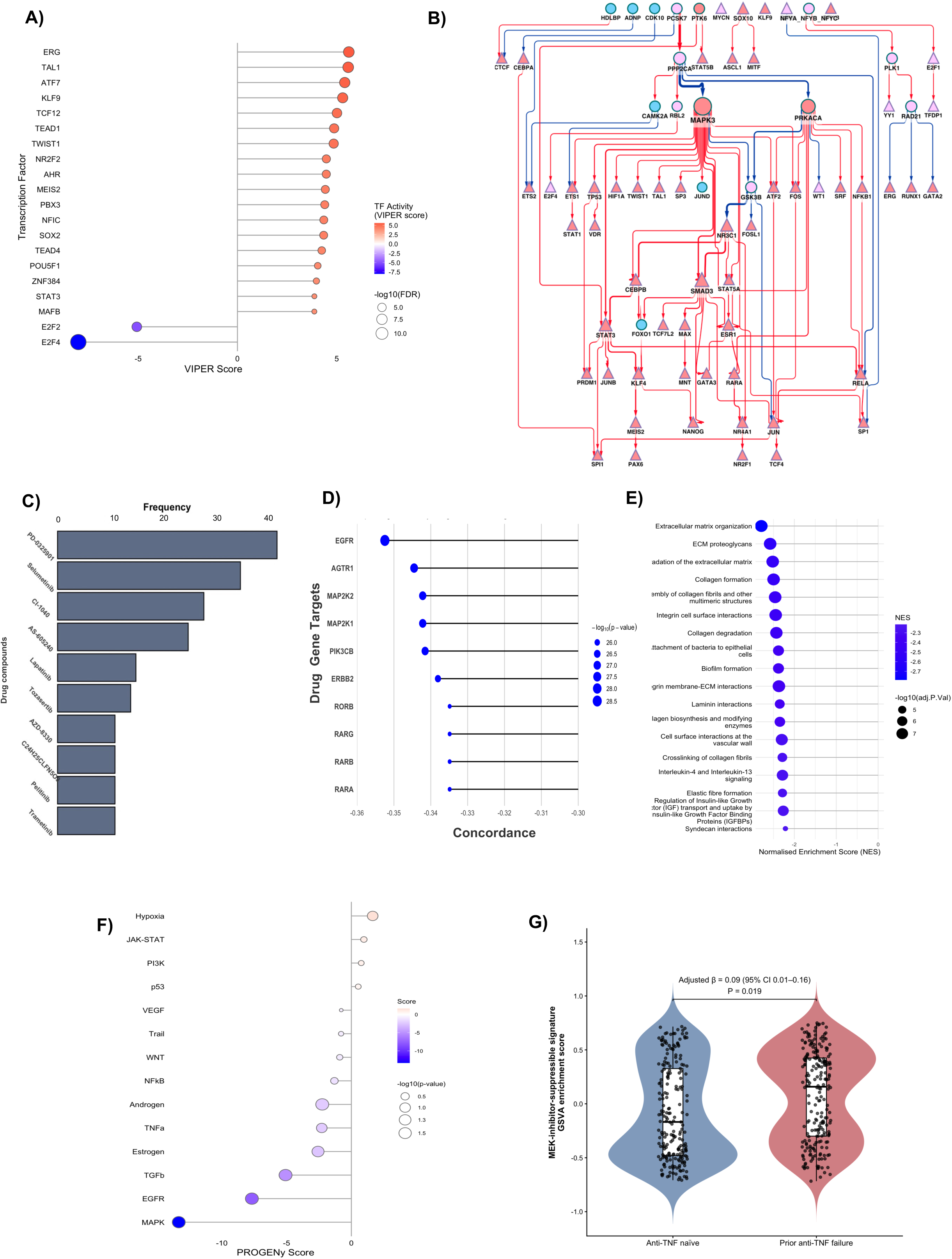
Regulatory-network and perturbational analyses prioritise MAPK/EGFR signalling as a key component of the UC anti-TNF-failure-associated molecular phenotype. **(A)** The 20 transcription factors with the largest absolute differences in inferred activity between prior anti-TNF-failure and anti-TNF-naïve groups, estimated using VIPER with DoRothEA regulons. Positive VIPER scores indicate higher inferred activity in patients with prior anti-TNF failure and negative scores indicate higher activity in anti-TNF-naïve patients. Point colour represents the VIPER score and point size represents −log₁₀(FDR). **(B)** High-confidence causal signalling network reconstructed using CARNIVAL from VIPER transcription-factor activities and directed protein-protein interactions from OmniPath. Only interactions assigned the maximum CARNIVAL edge weight of 100 are depicted. Node colour indicates inferred protein activity (red, increased; blue, reduced in prior anti-TNF failure), and node size is proportional to out-degree. Red and blue edges indicate positive and negative regulatory interactions, respectively, with edge width proportional to edge betweenness. MAPK3 occupies a highly connected upstream position within the inferred signalling network. **(C)** Frequency of the ten most recurrent drug compounds among negatively concordant LINCS L1000 perturbagen signatures predicted to reverse the anti-TNF-failure-associated expression profile. Counts incorporate signatures generated across the available cell lines, concentrations and exposure durations. **(D)** Gene targets ranked by negative concordance with the anti-TNF-failure-associated transcriptional profile. More negative weighted Pearson correlation coefficients indicate stronger predicted reversal; point size represents −log₁₀(*P* value). **(E)** Reactome gene-set enrichment analysis of paired UC colonic biopsy explants treated *ex vivo* with PD-0325901 versus vehicle. Selected significantly negatively enriched pathways are shown; negative NES values indicate suppression following MEK inhibition. Point size represents −log₁₀(Benjamini–Hochberg-adjusted *P* value), and colour represents the NES. **(F)** PROGENy-inferred pathway-activity changes following MEK inhibition in UC explants. Negative scores indicate reduced inferred activity following PD-0325901 treatment. Point colour represents the pathway-activity score and point size represents −log₁₀(*P* value). ECM, extracellular matrix; FDR, false-discovery rate; LINCS, Library of Integrated Network-based Cellular Signatures; MEK, mitogen-activated protein kinase kinase; NES, normalised enrichment score; TF, transcription factor; UC, ulcerative colitis. **(G)** Distribution of MEK-inhibitor-suppressible signature enrichment scores in baseline UNIFI colonic biopsies, stratified by prior anti-TNF status. The signature comprised the 100 most statistically significant genes downregulated following PD-0325901 treatment of UC explants, restricted to genes with Benjamini-Hochberg-adjusted *P* < 0.05. Higher GSVA scores indicate greater baseline enrichment of genes suppressed by MEK inhibition. Points represent individual participant scores. The displayed coefficient, 95% confidence interval and *P* value correspond to the association with prior anti-TNF failure from multivariable linear regression adjusted for log₂(CRP + 1), log₂(disease duration + 1) and total Geboes score. Anti-TNF-naïve patients constituted the reference group.

To functionally annotate the transcriptional regulators associated with prior anti-TNF failure, Gene Ontology (GO) over-representation analysis was performed on the regulons of transcription factors with significantly altered inferred activity. Several transcription factors with increased activity, including TEAD1, TCF4, RUNX2, MEF2A and KLF6, had regulons enriched for processes related to cell matrix adhesion, focal adhesion organisation, cytoskeletal remodelling, epithelial, endothelial and fibroblast migration, wound healing and extracellular-matrix organisation. These findings were consistent with the stromal and vascular enrichment identified by previous analyses.

Transcription factors with reduced inferred activity included IRF3 and FOXP1, whose regulons were enriched for cytokine responsive and lymphocyte differentiation programmes, and E2F2 and E2F4, whose regulons were strongly associated with DNA replication and cell cycle regulation. Conversely, increased STAT5B activity was associated with a regulon enriched for leukocyte and T-cell proliferation.

Collectively, these findings indicate that prior anti-TNF failure is associated with altered transcriptional regulatory activity involving stromal and tissue remodelling programmes, alongside suppression of specific immune pathways. This suggests that drug sequencing depreciation in UC may, at least in part, reflect upstream alterations in transcriptional regulatory networks that shape the mucosal inflammatory landscape in anti-TNF failure.

### Causal network inference prioritises MAPK3 as a central upstream signalling hub associated with anti-TNF failure

To investigate the upstream signalling architecture associated with transcriptional dysregulation in patients with prior anti-TNF failure, CARNIVAL causal network inference was applied^15^. This framework integrated transcription factor activities inferred using VIPER and curated directed regulatory interactions obtained from OmniPath. This enabled reconstruction of a context-specific network linking upstream signalling proteins with downstream transcriptional regulators.

The initial reconstructed network comprised approximately 6,000 nodes and 23,000 edges, representing proteins and regulatory interactions, respectively. To increase stringency and improve interpretability, the network was filtered to retain only interactions assigned the maximum confidence weight of 100. The resulting high-confidence network comprised 200 nodes and 250 edges (Figure 2B).

Network analysis identified several highly connected signalling proteins. MAPK3, which encodes ERK1, exhibited the highest out-degree centrality and was connected to multiple downstream signalling intermediates and transcription factors previously identified as dysregulated in patients with prior anti-TNF failure. Several MAPK3-associated interactions also displayed high edge betweenness, indicating that they lay along a large number of shortest paths connecting upstream signals with downstream regulatory programmes.

These network properties indicate MAPK3 is a central upstream signalling hub within the inferred signalling network associated with anti-TNF failure in UC. Its position provides a potential mechanistic link between the increased MAPK pathway activity identified in preceding analyses and the downstream transcriptional changes observed in anti-TNF failure. Collectively, these findings nominate MAPK3 as a candidate regulator of the molecular programme associated with prior anti-TNF failure.

### Connectivity mapping prioritises MAPK- and EGFR-targeted compounds following anti-TNF failure in UC

Given the substantial clinical challenge posed by drug sequencing depreciation following anti-TNF failure, we next sought to identify candidate therapeutic strategies that could potentially be repurposed for this difficult-to-treat patient population.

To achieve this, connectivity mapping was performed using more than 140,000 chemical perturbation signatures available through the NIH LINCS database^16^. The complete ranked differential gene expression results derived from anti-TNF-failed versus anti-TNF-naïve UC patients in the UNIFI cohort were used as the query signature. Compounds predicted to reverse the transcriptional changes associated with anti-TNF failure were identified on the basis of significant negative concordance with this gene expression profile.

This analysis identified 1,132 negatively concordant perturbagen signatures, indicating their predicted capacity to counteract the molecular phenotype associated with anti-TNF failure. Compounds targeting the MAPK signalling pathway emerged as the most prominent drug class, including the MEK inhibitors PD-0325901, selumetinib and CI-1040 (Figure 2C). Inhibitors of EGFR signalling, including the dual EGFR/ERBB2 inhibitor lapatinib, were also highly represented, consistent with the role of EGFR as an upstream activator of the MAPK signalling cascade.

Target-level analysis further supported these findings, with EGFR exhibiting the strongest negative concordance among the highest-ranked molecular targets (Figure 2D). AGTR1 was also highly ranked, followed by the MAPK pathway components MAP2K2 and MAP2K1, which encode MEK2 and MEK1, respectively. Other leading targets included PIK3CB and ERBB2. The prominence of the receptor tyrosine kinases EGFR and ERBB2 alongside the downstream MEK kinases MAP2K1 and MAP2K2 within the EGFR-MAPK pathways provided further support for the therapeutic relevance of this signalling axis.

To further prioritise therapeutically relevant candidates, analyses were subsequently restricted to perturbagen signatures generated in colonic cell lines. 91 significantly negatively concordant signatures met these criteria. The strongest negatively concordant perturbagen with an annotated molecular target was PD-98059 in the HT29 colonic epithelial cell line (weighted Pearson correlation = −0.31, *p* = 3.83 × 10⁻²²). PD-98059 prevents the activation of MAP2K1 (MEK1), thereby attenuating downstream ERK1/2 signalling - notably, MAPK3 (ERK1) was also identified as a central signalling hub in the CARNIVAL network analysis. Consistent with these network-level findings, MAPK pathway inhibitors constituted the dominant mechanistic class among the colonic cell line-derived perturbagens. Of the 91 significant signatures, 21 (23.1%) were generated by perturbagens targeting MAP2K1 and/or MAP2K2, substantially exceeding the representation of any other mechanistic class.

Taken together, these findings convergently nominate MAPK signalling as a central molecular pathway associated with anti-TNF failure and identify MEK/ERK inhibition as a candidate therapeutic strategy for further evaluation in anti-TNF-failed UC patients. The convergence of pathway activity analysis, causal network modelling and connectivity mapping on the MAPK axis provides a strong rationale for further functional evaluation of MAPK-targeted therapies in this patient subgroup.

### MEK inhibition suppresses molecular programmes associated with anti-TNF failure

To investigate whether the MAPK signalling axis contributes to or maintains the molecular phenotype associated with anti-TNF failure, we next examined whether pharmacological inhibition of this pathway could reverse the transcriptional programmes identified in anti-TNF-failed UC patients.

To address this question, transcriptomic data from colonic biopsy explants obtained from patients with active UC and treated *ex vivo* with the selective MEK inhibitor PD-0325901 were analysed. These samples were selected from a previously published dataset generated by Stankey and colleagues ^17^, in which paired biopsy explants were cultured with PD-0325901 or vehicle control for 18 hours before RNA sequencing

Differential expression and GSEA pathway analyses (Figure 2E) demonstrated that MEK inhibition markedly suppressed the stromal and extracellular matrix programmes found to be enriched in anti-TNF-failed patients, including pathways related to extracellular matrix organisation, collagen formation, and collagen degradation. In addition, MEK inhibition significantly reduced the activity of MAPK and EGFR signalling pathways as inferred through PROGENy analysis, both of which were identified as the most upregulated PROGENy signalling cascades in the anti-TNF-failed colonic mucosa (Figure 2F).

To capture the gene-level transcriptional response to MEK inhibition, a MEK-inhibitor-suppressible signature was defined using the 100 genes with the smallest Benjamini–Hochberg-adjusted *P* values among genes significantly downregulated following PD-0325901 treatment (adjusted *P* < 0.05). We next assessed whether this explant-derived signature was enriched in baseline UNIFI colonic biopsies. The signature was more highly enriched in patients with prior anti-TNF failure, and this association persisted after adjustment for CRP, disease duration and total Geboes score (adjusted β = 0.089, 95% CI 0.015–0.164; *P* = 0.019; Figure 2G). These findings link the anti-TNF-failure-associated mucosal phenotype to a transcriptional programme that is suppressed by MEK inhibition in human UC tissue.

Collectively, these findings therefore support a model in which MAPK-EGFR signalling acts upstream of the stromal remodelling programme associated with UC anti-TNF failure. Furthermore, these results provide functional support for the therapeutic prioritisation generated by connectivity mapping analyses, which identified MEK/ERK inhibitors among the most strongly negatively concordant perturbagens. Together, this suggests that pharmacological inhibition of the MAPK pathway may represent a potential strategy for reversing the molecular phenotype associated with anti-TNF failure in UC.

### Spatial localisation of MAPK/EGFR signalling identifies stromal-immune niches enriched for the anti-TNF failure molecular programme in inflamed UC mucosa

Having identified coordinated MAPK/EGFR signalling as a feature of the anti-TNF failure-associated molecular phenotype and shown that MEK inhibition modulated the corresponding stromal programmes in UC explants, we next sought to determine the cellular and spatial localisation of this signalling axis within inflamed UC mucosa. Single-cell-resolution spatial transcriptomic profiling was performed on FFPE colonic tissue sections from 16 patients with active UC and four non-IBD controls using the CosMx Spatial Molecular Imager (Supplementary Table 1).

A stromal-centred neighbourhood analysis was performed by defining a 50μm radius around each stromal index cell, including fibroblast subsets, endothelial cells and pericytes. Clustering the resulting neighbourhood-composition profiles identified ten distinct cellular niches (N1-N10), each comprising different proportions of immune, stromal and epithelial cell populations (Figure 3A).

**Figure 3:**
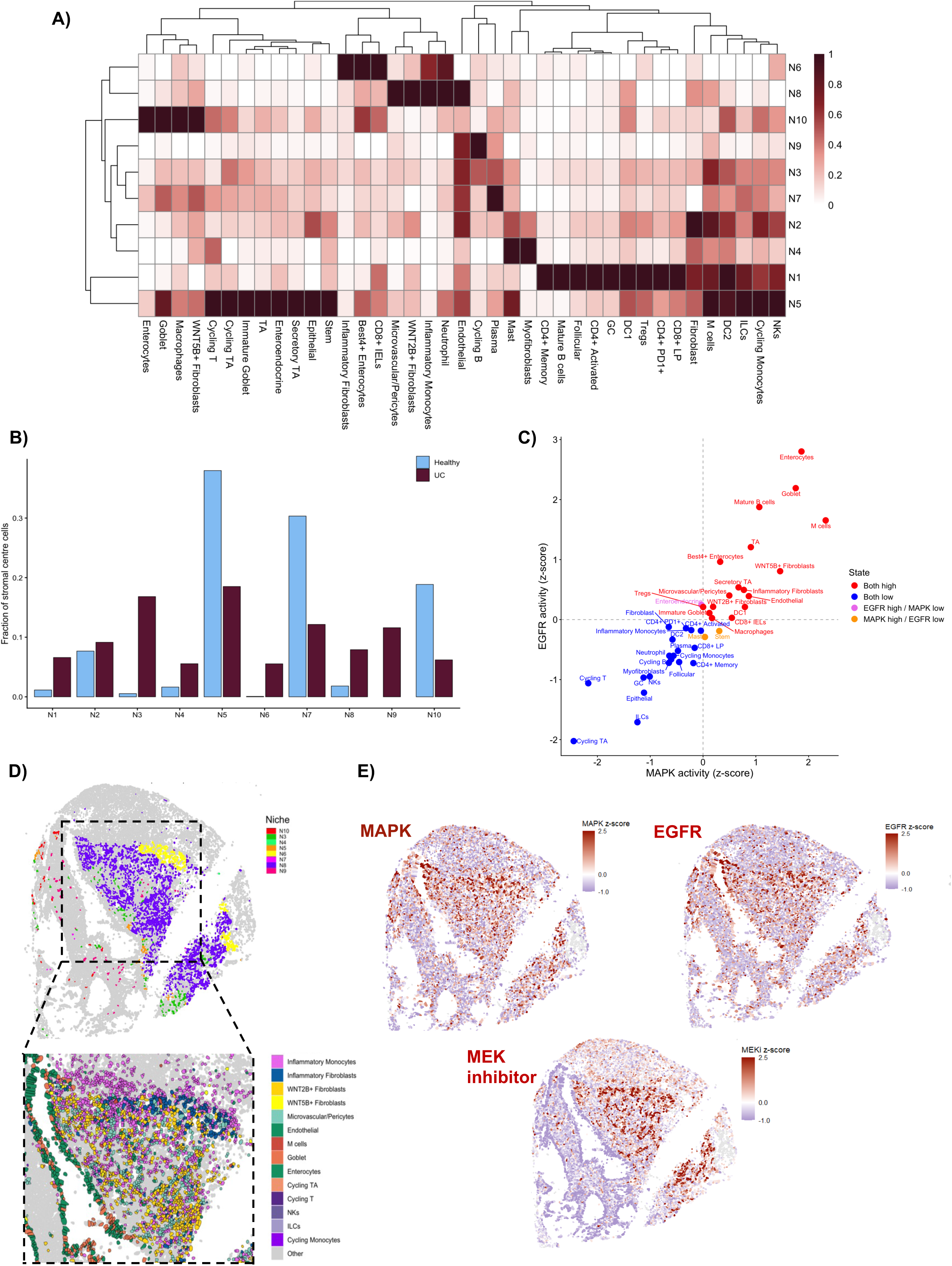
Single-cell-resolution spatial transcriptomics localises coordinated MAPK/EGFR activity and the MEK-inhibitor-suppressible programme to stromal-rich niches in inflamed UC mucosa. **(A)** Cellular composition of ten stromal-centred spatial niches identified by *k*-means clustering of cell-type proportions within 50µm neighbourhoods around fibroblast, endothelial and microvascular/pericyte index cells. Values represent mean cell-type proportions within each niche, min-max scaled separately for each cell type across niches from 0 (lowest relative representation) to 1 (highest relative representation). Consequently, colour intensity compares the distribution of each cell type across niches. Rows and columns were hierarchically clustered for visualisation. **(B)** Cohort-level pooled fraction of stromal index-cell neighbourhoods assigned to each niche in active UC and non-IBD control tissues. The non-IBD group is labelled “Healthy” in the panel. **(C)** Relationship between mean MAPK and EGFR pathway activities across annotated cell types. PROGENy scores were first averaged within each patient-cell-type combination and then across patients before being independently z-standardised across cell types. Each point represents one cell type. Dashed lines at zero divide cell types into those with relatively high or low activity; colours indicate concordantly high activity, concordantly low activity, MAPK-high/EGFR-low activity or EGFR-high/MAPK-low activity. **(D)** Spatial organisation of stromal-centred niches in a representative active-UC tissue section. The upper panel shows niche assignments of stromal index-cell neighbourhoods. The boxed region is magnified below to show the spatial proximity of selected fibroblast, vascular, myeloid, epithelial and lymphoid populations. **(E)** Spatial maps of inferred MAPK activity, inferred EGFR activity and the MEK-inhibitor-suppressible signature within the same representative UC tissue region. MAPK and EGFR activities were inferred using PROGENy. Each feature was independently z-standardised across cells within the displayed region; warmer colours indicate higher relative scores and cooler colours indicate lower relative scores. ILC, innate lymphoid cell; LP, lamina propria; MEK, mitogen-activated protein kinase kinase; TA, transit-amplifying; UC, ulcerative colitis.

N6 and N8 emerged as two distinct stromal-dominant niches. N6 was characterised principally by inflammatory fibroblasts alongside BEST4^+^ enterocytes, CD8^+^ intraepithelial lymphocytes, neutrophils, and inflammatory monocytes, whereas N8 contained prominent WNT2B⁺ fibroblast, inflammatory monocyte, microvascular/pericyte and endothelial populations. Together, these niches represented fibroblast- and vascular-rich stromal-myeloid microenvironments consistent with active inflammation and tissue remodelling.

The cohort-level frequency of these niches differed markedly between active UC and non-IBD control tissues (Figure 3B). N1, N3, N4, N6, N8 and N9 were more frequent in UC tissue, with N6 and N9 not detected in the non-IBD control samples. N8 was also substantially more frequent in UC than in non-IBD tissue. The enrichment of N6 and N8 in UC supported their identification as disease-associated stromal microenvironments.

Next, we evaluated the inferred activity of the MAPK and EGFR pathways at single-cell resolution across the spatial tissues. Across all cells, MAPK and EGFR pathway-activity scores were strongly positively correlated (Figure 3C). A subset of cells exhibited concurrently high inferred activity of both pathways. Notably, this subset comprised predominantly stromal cell types prominent within N6 and N8 neighbourhoods, including inflammatory fibroblasts, WNT5B⁺ fibroblasts, microvascular/pericyte cells and endothelial cells. Several epithelial populations, including enterocytes and goblet cells, also demonstrated high activity of both pathways. Conversely, most lymphoid populations, including cycling T cells, B cells and NK cells, exhibited low inferred activity of both pathways.

We also evaluated whether the transcriptional programme identified following MEK inhibition of UC explant cultures preferentially mapped to the stromal and vascular populations exhibiting coordinated MAPK and EGFR activity. Cell type-level enrichment analysis demonstrated that the MEK-inhibitor-suppressible signature was predominantly enriched within stromal and vascular compartments (Supplementary Figure 1). The strongest positive enrichment was observed in microvascular/pericyte and endothelial populations, both with z-scores greater than 3, followed by inflammatory fibroblasts and WNT5B⁺ fibroblasts. M cells demonstrated more modest positive enrichment, while myofibroblasts and WNT2B⁺ fibroblasts also exhibited positive scores. In contrast, most lymphoid and epithelial populations including mature B cells, enterocytes, goblet cells, enteroendocrine cells and stem/transit-amplifying populations demonstrated negative enrichment scores. These findings indicate that the top genes suppressed by MEK inhibition in whole-tissue explants predominantly reflects programmes expressed within the stromal and vascular compartments of colonic tissues.

The spatial architecture of N6 and N8 microenvironments was subsequently visualised in a representative UC tissue section (Figure 3D). This demonstrates the close spatial proximity of inflammatory monocytes, inflammatory fibroblasts, WNT2B⁺ and WNT5B⁺ fibroblasts, microvascular/pericyte cells and endothelial cells, together with interspersed epithelial and immune populations. These findings further illustrate the complex stromal-immune organisation of N6- and N8-enriched tissue regions.

Finally, we examined the spatial distribution of inferred MAPK activity, EGFR activity and the MEK-inhibitor-suppressible signature within the same representative UC tissue section. Regions with high inferred MAPK and EGFR activity showed substantial spatial concordance and coincided with areas containing prominent representation of the stromal-dominant N6 and N8 niches (Figure 3E). These regions also exhibited high enrichment scores for the MEK-inhibitor-suppressible signature, supporting the spatial convergence of coordinated MAPK/EGFR signalling and the explant-derived transcriptional programme within organised stromal-rich microenvironments. This spatial convergence identified the tissue regions in which coordinated MAPK/EGFR activity and the explant-derived MEK-inhibitor-suppressible programme were most strongly expressed.

Together, these findings indicate that the MAPK/EGFR-associated and MEK-inhibitor-suppressible component of the anti-TNF failure-associated molecular phenotype preferentially localises to organised stromal-immune niches within the inflamed UC mucosa. Microvascular/pericyte cells, endothelial cells and inflammatory fibroblast subsets represent the principal cellular compartments in which these programmes converge.

### Spatial ligand-receptor analysis identifies reciprocal stromal-myeloid communication within UC-enriched stromal niches

Having defined the cellular composition and signalling landscape of UC-associated stromal-immune niches, we next investigated the intercellular communication networks that may contribute to their organisation and maintenance. CellChat was used to infer ligand-receptor interactions among cells within the spatially adjacent N6 and N8 niches (Figure 4A) and to compare communication between UC and non-IBD tissues.

**Figure 4.**
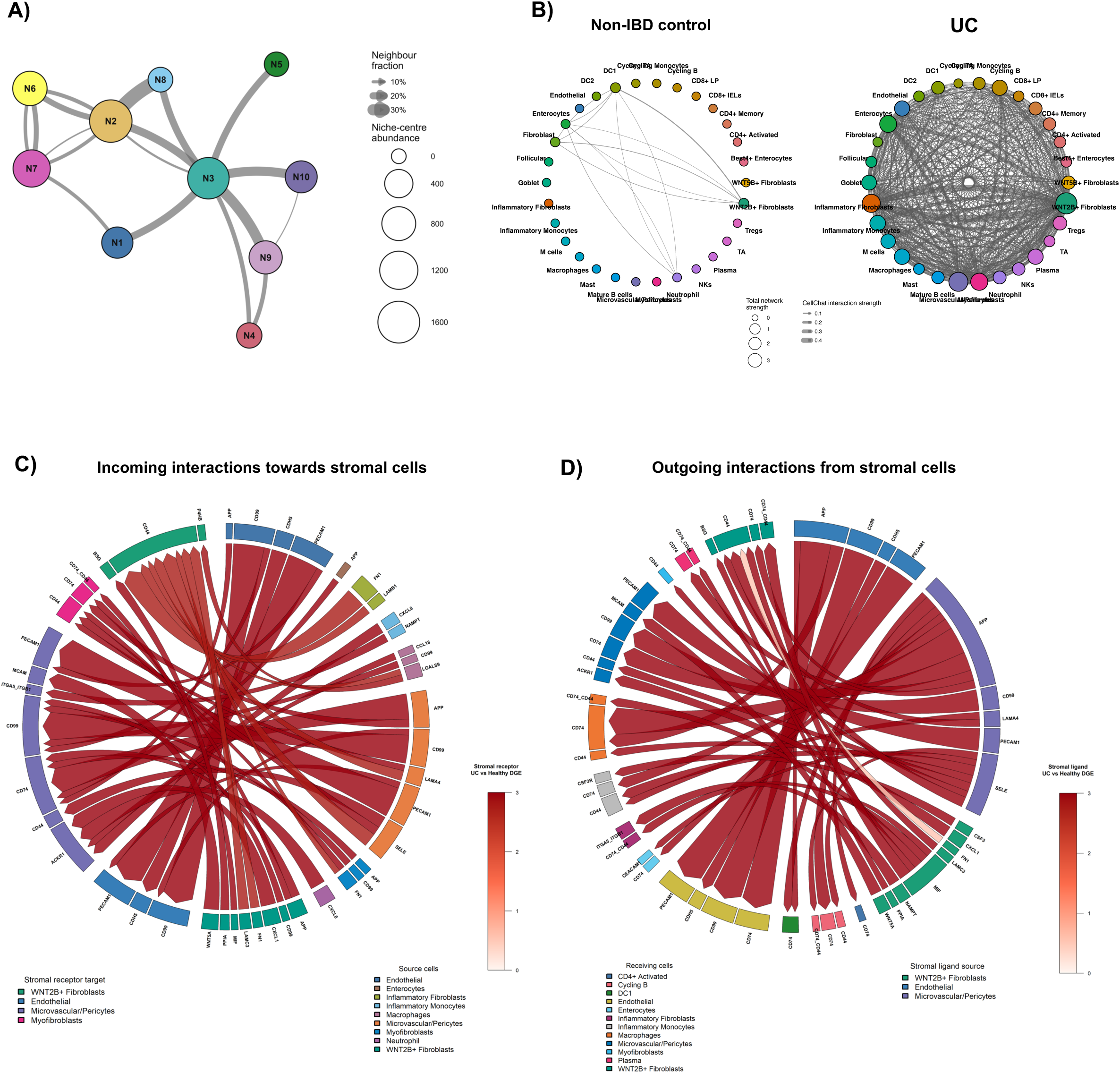
Spatial adjacency and ligand-receptor analysis identify reciprocal stromal-myeloid communication within UC-enriched stromal niches. **(A)** Spatial adjacency network of the ten stromal-centred niches in UC. Nodes represent niches N1–N10, node size indicates the abundance of niche-centre cells, and edge width represents the proportion of neighbouring niche centres belonging to the connected niche. **(B)** Global CellChat communication networks within the N6/N8-associated stromal compartment in non-IBD control and UC tissues. Nodes represent cell populations and are sized according to total network strength, calculated from combined incoming and outgoing communication. Directed edges represent predicted cell-cell communication, with edge width proportional to aggregate CellChat interaction strength. **(C)** Chord diagram showing UC-increased incoming ligand-receptor interactions received by stromal cell types. Ribbons connect ligands expressed by the indicated source populations to receptors expressed by stromal target populations. Ribbon width represents the increase in CellChat interaction probability in UC relative to non-IBD controls, while ribbon colour represents the UC-versus-control differential expression magnitude of the corresponding stromal receptor. **(D)** Chord diagram showing UC-increased outgoing ligand-receptor interactions produced by stromal cell types. Ribbons connect stromal ligands with receptors expressed by the indicated recipient populations. Ribbon width represents the increase in CellChat interaction probability in UC relative to non-IBD controls, while ribbon colour represents the UC-versus-control differential-expression magnitude of the corresponding stromal ligand. Only interactions meeting the CellChat significance criterion (log2FC>0.25, p adj <0.05) and exhibiting greater inferred communication probability in UC are displayed in panels C and D. Node and sector colours identify the corresponding niche or cell population. Abbreviations: DC, dendritic cell; IEL, intraepithelial lymphocyte; LP, lamina propria; TA, transit-amplifying; UC, ulcerative colitis.

Global CellChat adjacency analysis revealed a pronounced expansion in the breadth and strength of predicted intercellular communication within the UC N6–N8 compartment (Figure 4B). The non-IBD network contained relatively few interactions involving a restricted subset of cell populations, whereas UC tissues exhibited a densely interconnected communication network encompassing stromal, vascular, myeloid, epithelial and lymphoid populations. This global increase in network connectivity indicated marked reorganisation of intercellular communication within UC-associated stromal niches.

To characterise the direction and molecular composition of this communication, stromal populations were subsequently examined separately as signal-receiving and signal-producing populations. The analysis was restricted to interactions exhibiting greater inferred communication probability in UC than in non-IBD controls.

Analysis of incoming communication demonstrated extensive predicted signalling to inflammatory fibroblasts, WNT2B⁺ fibroblasts, endothelial cells, microvascular/pericyte cells and myofibroblasts from stromal, myeloid, epithelial and lymphoid populations (Figure 4C). These interactions included extracellular matrix and adhesion-associated ligands, such as FN1 and laminin components, together with mediators associated with inflammatory signalling, immune regulation, and chemotaxis, including MIF, NAMPT, LGALS9, CXCL8 and CCL18. In particular, predicted signals originating from inflammatory monocytes, macrophages and other myeloid populations converged on fibroblast and vascular cell populations, indicating that stromal cells within N6 and N8 integrate incoming inflammatory signals from the surrounding myeloid compartment alongside matrix- and adhesion-associated inputs.

Outgoing communication further highlighted a prominent stromal-myeloid signalling axis within the combined N6-N8 microenvironment in UC compared to control tissues (Figure 4D). WNT2B⁺ and inflammatory fibroblasts, endothelial cells, microvascular/pericyte cells and myofibroblasts generated predicted signals towards inflammatory monocytes, macrophages and other myeloid recipients. Upregulated interactions in UC included CXCL1-CXCR1/2 and CSF3-CSF3R signalling, consistent with stromal regulation of myeloid recruitment and activation, together with MIF-CD74/CD44 and NAMPT-associated inflammatory signalling. Additional WNT5A-FZD, APP, FN1 and laminin-associated interactions mediated communication between stromal, vascular, epithelial and immune populations.

Thus, stromal populations within the adjacent N6 and N8 niches acted as both recipients and producers of increased communication signals in UC. The combination of incoming myeloid-to-stromal and outgoing stromal-to-myeloid interactions supports the presence of a reciprocal stromal-myeloid signalling circuit linking the fibroblast- and vascular-rich architecture of N8 with the myeloid-enriched composition of N6. These findings suggest that reciprocal stromal-myeloid crosstalk may contribute to the organisation and maintenance of UC-enriched tissue microenvironments that localise the molecular programme associated with prior anti-TNF failure.

## DISCUSSION

Patients with prior anti-TNF failure representing a clinically challenging subgroup of UC. They experience poorer outcomes with several subsequent advanced therapies, yet the biological basis of this treatment-refractory state remains poorly understood^18^. In this study, we integrated clinical, transcriptomic, perturbational and spatial analyses to define the molecular phenotype associated with prior anti-TNF failure in UC.

Using data from the UNIFI phase III trial, we found that patients who had been previously exposed to anti-TNF therapies had a greater baseline disease burden compared to those who were anti-TNF naïve. Prior anti-TNF failure in ustekinumab-treated participants was associated with lower odds of achieving clinical response and mucosal healing compared to anti-TNF naïve patients, even after adjusting for baseline differences in disease burden, namely CRP, disease duration, and histological activity. Endoscopic healing and clinical remission showed directionally consistent associations that did not reach statistical significance. These findings suggest that subsequent poor treatment outcomes in anti-TNF failed patients, we term drug-sequencing depreciation, is not fully explained by conventional clinical measures of disease severity and indicates there may be a distinct molecular phenotype underpinning these patients.

Multimodal transcriptomic evaluation revealed that the molecular phenotype associated with prior anti-TNF failure was dominated by stromal activation and extracellular matrix (ECM) remodelling. Despite relatively modest differences in individual gene expression, pathway-level analyses identified coordinated enrichment of ECM organisation, collagen remodelling and integrin-associated programmes after adjustment for measured differences in clinical markers of disease severity. Cellular deconvolution indicated increased representation of stromal cell types such as fibroblasts, endothelial cells and pericytes in anti-TNF failed UC mucosa, accompanied by relative reductions in selected T-cell populations. Transcription factor activity inference further identified regulators associated with cell-matrix adhesion, migration, wound healing and ECM organisation. Together, these findings indicate that treatment refractoriness in UC is associated with dysregulation of the stromal compartment.

Multiple independent analyses also converged on MAPK/EGFR signalling as a prominent component of this phenotype. Pathway activity inference identified increased MAPK, EGFR, PI3K and TGF-β activity in anti-TNF-failed mucosa, whereas activity of TNF, NF-κB and JAK-STAT pathways was relatively reduced. This suggests a relative shift towards growth factor, repair and remodelling pathways following anti-TNF failure. Causal network reconstruction prioritised MAPK3 as a highly connected upstream signalling hub, while connectivity mapping independently identified MEK and EGFR inhibitors among the perturbations most strongly predicted to reverse the anti-TNF failure-associated transcriptional phenotype.

The predicted therapeutic vulnerability was supported using an independent transcriptomics dataset of human UC biopsy explant cultures treated *ex vivo* with the MEK inhibitor PD-0325901^17^. MEK inhibition suppressed ECM organisation and collagen remodelling programmes and reduced inferred MAPK and EGFR activity. These findings demonstrate that transcriptional programmes associated with prior anti-TNF failure are suppressible through MAPK-pathway perturbation in human inflamed colonic tissue.

Single cell-resolution spatial profiling provided anatomical context for these findings. Spatial neighbourhood analysis revealed stromal-immune enriched tissue niches (N6 and N8) in UC compared to non-IBD colonic tissues, containing inflammatory and WNT-associated fibroblasts, inflammatory monocytes, neutrophils, endothelial cells and microvascular/pericyte populations. Mean MAPK and EGFR pathway activities were strongly correlated across cell types and were highest in stromal populations prominent within the N6 and N8 niches. A MEK-inhibitor-suppressible signature similarly mapped most strongly to microvascular/pericyte cells, endothelial cells and inflammatory fibroblast subsets. These results localise the implicated programme to spatially organised stromal-immune microenvironments rather than to a single cellular lineage.

Spatial ligand-receptor inference further suggested that N6 and N8 are linked through reciprocal stromal-myeloid communication. Predicted myeloid-to-stromal signals included MIF, CXCL8 and CCL18, whereas stromal outgoing interactions included CXCL1-CXCR1/2 and CSF3–CSF3R signalling towards myeloid populations. MIF-CD74/CD44 signalling also provides a plausible connection between stromal-myeloid cross-talk and MAPK/ERK activity^19,20^. Although these interactions require experimental validation, they support a model in which reciprocal signalling between fibroblast-rich and myeloid-rich niches helps sustain an inflammatory and tissue-remodelling microenvironment.

Previously thought to be structural support cells, these observations align with mounting evidence that stromal cells may actively sustain intestinal inflammation through effects on immune cell recruitment, cytokine release, angiogenesis, epithelial repair and tissue remodelling^21^. Inflammatory fibroblast and myeloid cell programmes have previously been associated with anti-TNF non-response in IBD^22,23^. Our findings extend this concept by identifying a spatially organised stromal phenotype associated with anti-TNF failure in UC.

As the spatial cohort was not stratified by previous anti-TNF failure, however, these data localise an independently derived anti-TNF-failure-associated programme within inflamed UC tissue but do not independently demonstrate that these niches are selectively enriched in anti-TNF-failed patients. Furthermore, whether this stromal state precedes anti-TNF exposure or is progressively reinforced by unsuccessful treatment remains unresolved. A recent longitudinal single-cell transcriptomics study reported increased fibroblast activation and neutrophil-recruiting programmes following anti-TNF non-response^24^, suggesting that ineffective therapy and persistent inflammation may consolidate pre-existing stromal circuits. Previous studies, however, have also indicated that stromal programmes may portend treatment resistance^25,26^. Prospective sampling before and after treatment will be required to distinguish a baseline resistance endotype from a molecular state acquired or amplified following therapeutic exposure. Furthermore, whilst our findings indicate that MEK inhibitors may be an attractive therapeutic candidate to reverse this molecular phenotype, they do not establish that MEK inhibition would be clinically effective in anti-TNF-failed UC. MAPK/EGFR signalling also contributes to epithelial restitution and tissue homeostasis, and broad systemic inhibition could disrupt protective repair processes^27,28^. Therefore, therapeutic exploitation of this signalling axis may require selective targeting of pathogenic stromal or myeloid populations rather than global pathway inhibition. Finally, anti-TNF failure encompasses heterogeneous pharmacokinetic, immunogenic and mechanistic causes which have not been resolved in this study, and several cellular and pathway-level findings remain computational inferences.

Nevertheless, this study is strengthened by the use of a large, well-characterised clinical trial cohort, adjustment for measured differences in disease burden, and convergence across transcriptomic, regulatory network, perturbational, *ex vivo* and spatial analyses. Orthogonal protein-level and structural validation, together with prospective clinical studies and *in vivo* functional work, will be required to establish whether this phenotype predicts treatment resistance and whether it can be safely targeted.

In conclusion, prior anti-TNF failure in UC is associated with a treatment-refractory state characterised by stromal and vascular expansion, ECM remodelling and a relative shift towards MAPK/EGFR signalling. Human explant and spatial analyses support the biological relevance and tissue localisation of this programme and nominate the MAPK-ERK axis for further therapeutic investigation. These findings highlight stromal biology as an important component of drug-sequencing depreciation and support the development of molecularly informed approaches to therapeutic sequencing in UC.

## METHODS

### Differential gene expression analysis of GSE206285

GEO Series GSE206285^9^ contains array-based transcriptomic data generated from baseline colonic biopsies collected during the UNIFI randomised, placebo-controlled phase III trial of ustekinumab in moderate-to-severe ulcerative colitis^8^. Robust multi-array average (RMA)-normalised expression profiles and associated participant metadata were downloaded from the NCBI Gene Expression Omnibus. Probe sets mapping uniquely to a single Entrez Gene identifier were retained. Where multiple probe sets mapped to the same gene, their expression values were summarised to obtain one gene-level measurement per sample.

Expression profiles were matched to clinical metadata using sample identifiers. Genes with an expression intensity greater than 4 in at least three samples were retained. Samples missing anti-TNF exposure status or any prespecified covariate were excluded by complete-case analysis. Prior anti-TNF status was encoded as a categorical variable, with anti-TNF-naïve patients serving as the reference group.

Gene-wise linear models were fitted using the limma package^29^. The model included prior anti-TNF failure as the exposure of interest and adjusted for baseline disease duration, Mayo Endoscopic Score, faecal calprotectin, serum C-reactive protein and total Geboes histological score. Empirical Bayes moderation was applied to the gene-wise variance estimates. The coefficient for prior anti-TNF failure represented the adjusted expression difference between patients with prior anti-TNF failure and anti-TNF-naïve patients. Positive log₂ fold changes therefore indicated higher expression in the prior anti-TNF-failure group. Multiple-testing correction was performed using the Benjamini-Hochberg method. For visualisation in volcano plots, genes with an absolute log₂ fold change greater than 0.5 and a false-discovery rate below 0.05 were classified as differentially expressed.

### Baseline clinical comparisons

Baseline clinical characteristics were compared between patients with prior anti-TNF failure and anti-TNF-naïve patients using two-sided Mann–Whitney U tests. Continuous variables, including disease duration, serum CRP and total Geboes histological score, were summarised using medians and interquartile ranges.

### Associations with ustekinumab induction outcomes

Among ustekinumab-treated participants, associations between prior anti-TNF failure and week 8 clinical response, clinical remission, endoscopic healing and mucosal healing were assessed using separate multivariable logistic regression models. Models were adjusted for baseline CRP, disease duration and total Geboes score. CRP and disease duration were transformed as log₂(x + 1). Complete-case analysis included 318 participants. Adjusted odds ratios and Wald 95% confidence intervals were reported, with anti-TNF-naïve patients as the reference group. Multicollinearity was assessed using variance inflation factors.

### Gene set enrichment analysis

Ranked gene-set enrichment analysis was performed using clusterProfiler^30^ and ReactomePA^31^. Genes were ranked in decreasing order according to the adjusted-model log₂ fold change associated with prior anti-TNF failure. Gene identifiers were mapped to Entrez Gene identifiers using org.Hs.eg.db, and genes without a valid mapping were excluded. Gene sets containing between 10 and 500 genes were included. Multiple-testing correction was performed using the Benjamini-Hochberg method, and pathways with an adjusted P value below 0.05 were considered significantly enriched. A positive normalised enrichment score indicated enrichment in patients with prior anti-TNF failure, whereas a negative score indicated enrichment in anti-TNF-naïve patients.

### Cellular deconvolution analysis

Relative cell-type enrichment was inferred from the UNIFI microarray profiles using xCell version 1.1.0^11^. The xCellAnalysis function was applied with rnaseq = FALSE, and analysis was restricted to prespecified cell populations normally represented in intestinal tissue. Composite stromal and immune scores were calculated by averaging the enrichment scores of the corresponding prespecified cell populations. Median scores were reported for the prior anti-TNF-failure and anti-TNF-naïve groups. Between-group differences were assessed using two-sided Mann-Whitney U tests, with Benjamini-Hochberg correction across the tested cell types.

As a complementary analysis focused on immune-cell composition, CIBERSORT^12^ was applied using the LM22 reference signature and default parameters. Estimated immune-cell fractions were compared between prior anti-TNF-failure and anti-TNF-naïve groups using two-sided Mann–Whitney U tests followed by Benjamini-Hochberg correction.

### Pathway activity inference

Activity of canonical signalling pathways was inferred using the PROGENy model implemented through decoupleR^14^. The moderated gene-level statistics from the differential-expression analysis were integrated with the signed and weighted PROGENy pathway–target network using the weighted-mean method. Pathways represented by fewer than five measured target genes were excluded, and significance was estimated using 100 permutations. Normalised weighted-mean scores were used as the pathway-activity estimates. Positive scores indicated greater inferred pathway activity in patients with prior anti-TNF failure, whereas negative scores indicated greater activity in anti-TNF-naïve patients.

For selected pathways, including MAPK, PI3K, TGFβ, TNFα and NF-κB signalling, the direction of differential expression of individual pathway-responsive genes was compared with the corresponding signed PROGENy network weights to illustrate the genes contributing to the inferred activity score. P values were adjusted across the evaluated pathways using the Benjamini-Hochberg method.

### Transcription factor activity and regulon enrichment

Directed, signed transcription factor-target interactions with DoRothEA confidence levels A–C were retrieved using OmniPathR^32^. Transcription-factor activity was inferred from limma moderated t-statistics using VIPER implemented through decoupleR. Regulons containing fewer than five measured targets were excluded, and pleiotropy correction was enabled. Positive scores indicated greater inferred activity in patients with prior anti-TNF failure. P values were adjusted using the Benjamini–Hochberg method.

Gene Ontology Biological Process over-representation analysis was performed on the target genes of each significant transcription factor using enrichGO in clusterProfiler. The background comprised all DoRothEA target genes represented in the expression dataset. Regulon functions with a false-discovery rate below 0.05 were considered significant.

### Upstream signalling network inference

Upstream signalling networks were inferred using inverse CARNIVAL^15^. VIPER transcription factor activity scores were supplied as measurements and directed protein-protein interactions retrieved from OmniPath formed the prior-knowledge network. runInverseCarnival was applied using default parameters and the IBM ILOG CPLEX optimisation solver. The initial solution contained 6,105 nodes and 23,642 directed interactions. Restriction to interactions with the maximum solution weight of 100 produced a high-confidence network containing 227 nodes and 254 edges. These interactions were retained for visualisation in Cytoscape. Node colour represented inferred protein activity, node size represented out-degree, edge colour indicated interaction sign and edge width represented edge betweenness.

### Connectivity mapping

Pharmacological perturbations predicted to reverse the anti-TNF-failure-associated transcriptional phenotype were identified using the iLINCS platform and its Library of Integrated Network-based Cellular Signatures L1000 resource^33^. The adjusted differential-expression signature comparing prior anti-TNF failure with anti-TNF-naïve UC was used as the query. Weighted Pearson connectivity scores were calculated against chemical-perturbagen signatures across available cell lines, concentrations and exposure durations. Perturbations showing negative concordance with the query signature at nominal P < 0.05 were considered candidate phenotype-reversing compounds and were subsequently restricted to signatures generated in colon-derived cell lines.

### Analysis of MEK-inhibitor-treated UC explants

Raw RNA-sequencing counts from paired UC biopsy explants treated for 18 hours with the selective MEK inhibitor PD-0325901 or vehicle were obtained from a previously published dataset^17,34^. Lowly expressed genes were removed using filterByExpr in edgeR^34^, followed by trimmed mean of M-values normalisation. Counts were transformed using the voom method, and gene-wise linear models were fitted using limma^29^. The design included donor and treatment, thereby accounting for the paired experimental design. Empirical Bayes moderation was applied, and the MEK-inhibitor effect was estimated relative to vehicle treatment. P values were adjusted using the Benjamini–Hochberg method.

Ensembl identifiers were mapped to HGNC gene symbols and Entrez Gene identifiers using org.Hs.eg.db. Reactome GSEA was performed using genes ranked by the MEK-inhibitor-associated log₂ fold change. Where multiple genes mapped to the same Entrez identifier, the entry with the largest absolute log₂ fold change was retained. Pathways with an adjusted P value below 0.05 were considered significant.

Changes in canonical pathway activity following MEK inhibition were inferred using PROGENy. Limma moderated *t*-statistics were integrated with the signed PROGENy network containing the 100 highest-weighted responsive genes per pathway using the weighted-mean method, with 100 permutations and a minimum regulon size of five genes. Negative activity scores indicated reduced inferred pathway activity following MEK inhibition.

The MEK-inhibitor-suppressible signature comprised the 100 genes with the smallest Benjamini–Hochberg-adjusted *P* values among genes significantly downregulated following MEK inhibition (adjusted *P* < 0.05). Per-sample signature enrichment scores were calculated in baseline UNIFI colonic biopsies using gene-set variation analysis. The association between prior anti-TNF failure and signature enrichment was assessed using multivariable linear regression, with the GSVA enrichment score as the dependent variable and adjustment for log₂(CRP + 1), log₂(disease duration + 1) and total Geboes score. Anti-TNF-naïve patients constituted the reference group.

### Single-cell-resolution spatial transcriptomics

#### Tissue preparation and spatial profiling

CosMx Spatial Molecular Imaging was performed on FFPE colonic biopsies from 16 patients with active UC and four non-IBD controls. Samples were obtained through the THAMES-IBD study under approval from the Yorkshire & The Humber–Sheffield Research Ethics Committee (22/YH/0043). Tissue integrity and pathological representation were assessed by haematoxylin and eosin staining before samples were selected for spatial profiling.

Five-micrometre sections were mounted onto Leica Bond Plus slides and incubated overnight at 60°C. Sections were deparaffinised using xylene and graded ethanol, followed by heat-mediated target retrieval at 100°C for 15 minutes and permeabilisation with Proteinase K (3 μg/mL) for 30 minutes at 40°C. After washing with 2× SSC-T and DEPC-treated water, fiducial markers were applied for image registration. Sections were subsequently post-fixed in 10% neutral-buffered formalin and treated with Sulfo-NHS-Acetate to minimise non-specific binding.

RNA targets were hybridised for 16-18 hours at 37°C using the CosMx Human Universal Cell Characterization Panel, comprising approximately 6,000 transcripts. Following stringent washing with SSC and formamide, nuclei were labelled with DAPI and cell boundaries identified using PanCK, CD45, CD3 and CD298/B2M segmentation markers. Slides were imaged using the CosMx Spatial Molecular Imager with manufacturer-recommended pre-bleaching, imaging and segmentation settings.

#### Primary processing and quality control

Raw imaging data were processed using the CosMx/AtoMx analysis pipeline. Cell boundaries were identified using Cellpose-based segmentation, and detected transcripts were assigned to individual cells. Cell-level expression matrices, metadata and spatial coordinates were subsequently exported for analysis in R.

Quality control was undertaken at both cell and field-of-view levels. Cell-level metrics included cell area, numbers of detected transcripts and genes, and the proportion of signal attributable to negative-control probes. Field-of-view quality was assessed using fluidic alignment, transcript localisation and transcript-assignment efficiency. Fields of view with a mean unassigned-transcript fraction greater than 20% were excluded.

#### Normalisation and dimensionality reduction

Downstream analysis was performed using Seurat. Expression data were normalised using SCTransform using Seurat^35^ and integrated across samples. Dimensionality reduction was subsequently performed using uniform manifold approximation and projection with a cosine-distance metric.

#### Stromal-centred neighbourhood analysis

Spatial neighbourhoods were defined around stromal and vascular index cells, comprising inflammatory, WNT2B⁺ and WNT5B⁺ fibroblasts, other fibroblasts, myofibroblasts, endothelial cells and microvascular/pericyte populations. Analysis was restricted to UC and non-IBD control tissues.

For each index cell, all annotated cells located within a 50-μm radius were identified within the same slide and field of view using global spatial coordinates and a conversion factor of 0.12028 μm per pixel. Each neighbourhood was represented by the proportional abundance of every annotated cell type within this radius. Neighbourhood-composition profiles were partitioned into ten niches using k-means clustering with 20 random initialisations and a fixed random seed. Niches containing fewer than 50 index-cell neighbourhoods were excluded.

The cellular composition of each niche was summarised by averaging cell-type proportions across its constituent neighbourhoods. For heatmap visualisation, values were independently rescaled between 0 and 1 for each cell type. Cell populations contributing less than 1% of a niche were combined into a low-abundance category in compositional plots. Niche frequencies were calculated as the proportion of stromal-centred neighbourhoods assigned to each niche within each disease group.

#### Cell type pathway and perturbation-signature analysis

Cell-level MAPK and EGFR activities were inferred from SCTransform-normalised expression data using PROGENy and the 100 highest-weighted responsive genes per pathway. To reduce bias from unequal cell numbers, scores were first averaged within each patient–cell-type combination and subsequently across patients. Mean MAPK and EGFR scores were independently standardised across cell types. Cell types were classified as having relatively high or low activity according to whether their standardised score was above or below the across-cell-type mean. Concordance between MAPK and EGFR activity was assessed across cell-type means using Pearson correlation.

The MEK-inhibitor-suppressible signature comprised genes downregulated following MEK inhibition in UC explant cultures. Cell-level signature scores were averaged within each patient–cell-type combination and subsequently across patients. Empirical P values were estimated using 2,000 permutations of cell-type labels and adjusted across cell types using the Benjamini-Hochberg method. Mean scores were z-standardised across cell types for visualisation.

#### Spatial mapping of pathway activity and the MEK-inhibitor-suppressible signature

For spatial visualisation, a representative UC tissue region comprising nine contiguous fields of view was selected. MAPK and EGFR activities were inferred in individual cells from SCTransform-normalised expression data using PROGENy and the 100 highest-weighted responsive genes per pathway. Scores were independently z-standardised across cells within the selected tissue region.

The MEK-inhibitor-suppressible signature comprised the 100 most statistically significant genes downregulated following MEK inhibition in UC explant cultures. Signature genes were intersected with genes represented in the CosMx dataset, and per-cell module scores were calculated from SCTransform-normalised expression data using Seurat’s AddModuleScore. This method calculates the average expression of signature genes relative to expression-matched control genes. Module scores were z-standardised across cells within the representative region for visualisation.

Cell-segmentation polygons were reconstructed from global boundary coordinates and joined to pathway and signature scores using unique cell identifiers. Spatial distributions of MAPK activity, EGFR activity and the MEK-inhibitor-suppressible signature were visualised alongside annotated cell populations in the same tissue region.

#### Spatial ligand-receptor analysis

To investigate cell–cell communication within stromal-associated spatial niches, cells assigned to N6 or N8 were combined, and stromal cells within this compartment were designated as spatial anchors. Within each tissue section, RANN was used to identify all cells located within 50 µm of at least one stromal anchor. Proximity searches were performed separately for each tissue section to prevent neighbourhoods spanning different samples.

CellChat^36^ was applied independently to cells pooled from non-IBD and UC tissues using CellChatDB.human. Spatially constrained communication probabilities were calculated using a truncated mean (trim = 0.1), a 50 µm interaction range, a 10 µm contact-dependent interaction range and 100 bootstrap iterations.

Ligand-receptor pairs were classified as UC-increased when they met the CellChat significance criterion in UC (*P* ≤ 0.05) and had a higher inferred communication probability in UC than in non-IBD controls (ΔP = P_UC − P_Healthy > 0). Outgoing communication was defined as stromal ligands interacting with receptors on neighbouring cells, whereas incoming communication comprised ligands expressed by neighbouring populations interacting with stromal receptors.

For visualisation in the chord diagrams, interactions were additionally required to have a corresponding stromal ligand, for outgoing communication, or stromal receptor, for incoming communication, with a UC-versus-control log₂ fold change >0.25 and a Benjamini–Hochberg-adjusted *P* value <0.05. Ribbon width represented the UC-control difference in CellChat communication probability, while ribbon colour represented the log₂ fold change of the corresponding stromal ligand or receptor.

Condition-specific global communication networks were visualised using netVisual_circle, with node size representing total incoming and outgoing network strength and edge width representing aggregate CellChat interaction strength. CellChat objects were combined using mergeCellChat, and overall interaction number and strength were compared using compareInteractions. Differential network structure and pathway-level information flow were additionally assessed using netVisual_diffInteraction and rankNet, respectively.

### Software

Analyses were performed in R version 4.5.2. Principal packages included limma 3.66.0, edgeR 4.8.2, AnnotationDbi 1.72.0, org.Hs.eg.db 3.22.0, clusterProfiler 4.18.4, ReactomePA 1.54.0, xCell 1.1.0, OmniPathR 3.10.1, decoupleR 2.16.0, viper 1.44.0, CARNIVAL 2.12.0, GSVA 2.4.7 and car 3.1-5. Spatial analyses used Seurat 5.4.0, SeuratObject 5.3.0, sctransform 0.4.3, progeny 1.32.0, sf 1.1.0, dbscan 1.2.4 and pheatmap 1.0.13. Mixed-effects modelling and estimated marginal means used lmerTest 3.2.1 and emmeans 2.0.3, respectively. Network visualisation was performed using Cytoscape version 3.10.1.

---

**Supplementary Figure 1.**
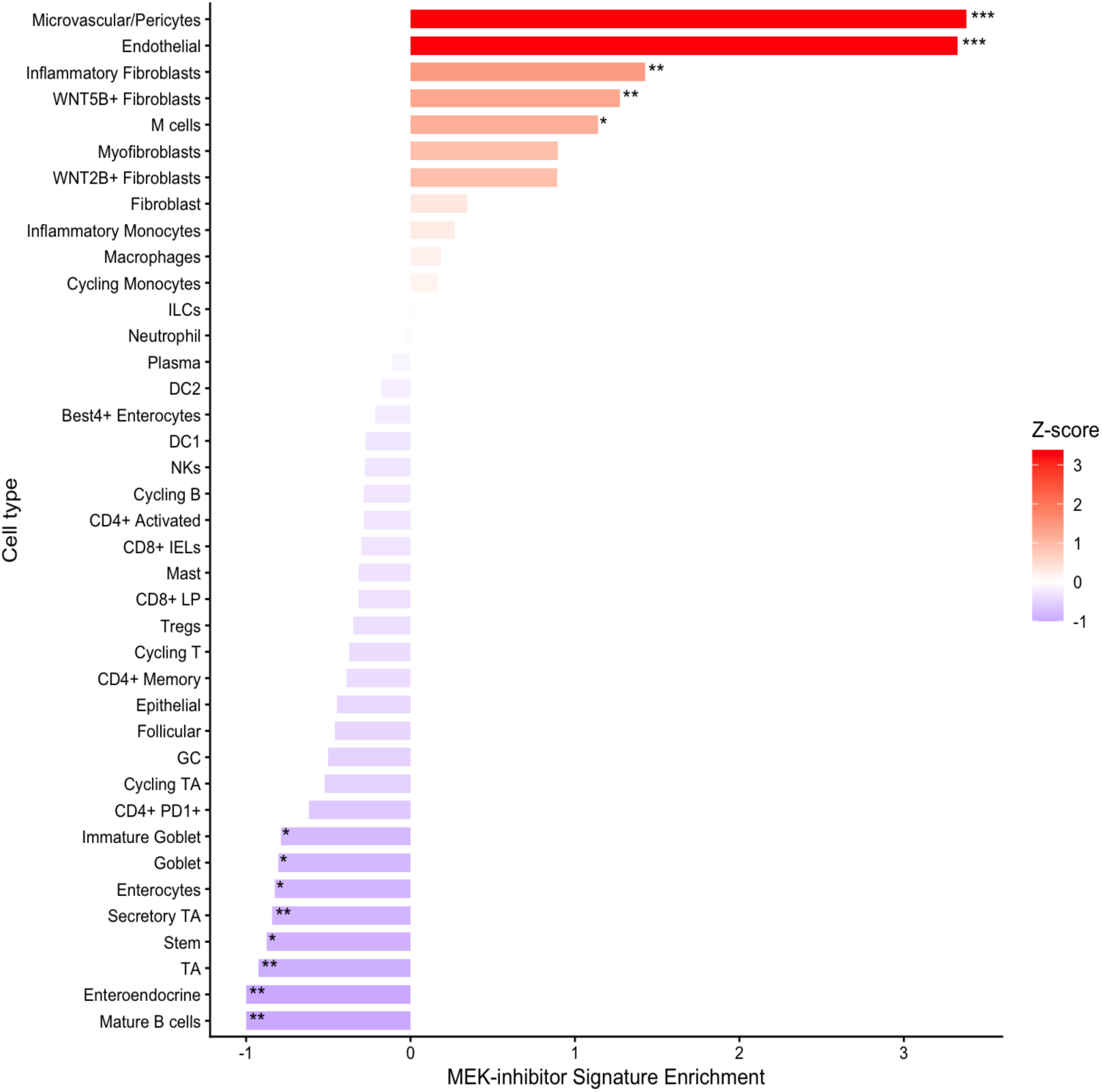
Cell-type distribution of the MEK-inhibitor-suppressible transcriptional signature in the spatial transcriptomic dataset. The signature comprised the 100 most statistically significant genes downregulated following PD-0325901 treatment of UC colonic biopsy explants, restricted to genes with Benjamini-Hochberg-adjusted *P* < 0.05. Signature scores were averaged within each patient-cell-type combination and subsequently across patients before being z-standardised across annotated cell types. Positive values therefore indicate relatively greater baseline expression of genes suppressed by MEK inhibition, whereas negative values indicate relatively lower expression. Bar colour represents the z-score. Empirical *P* values were estimated using 2,000 permutations of cell-type labels and adjusted across cell types using the Benjamini–Hochberg method. *Adjusted *P* < 0.05, **adjusted *P* < 0.01 and ***adjusted *P* < 0.001. ILC, innate lymphoid cell; LP, lamina propria; MEK, mitogen-activated protein kinase kinase; TA, transit-amplifying.

**Supplementary Table 1.** Clinical characteristics of the spatial transcriptomic cohort.

| Characteristic | Active UC cohort (N = 16) | Non-IBD control cohort (N = 4) |
| --- | --- | --- |
| Age at endoscopy, years | 34.5 (27.0-50.8) | 26.0 (21.5–33.0) |
| Female sex | 16 (100%) | 4 (100%) |
| Disease duration, years | 5.0 (2.0–21.5) | N/A |
| Disease extent |  |  |
| — Proctitis (E1) | 1 (6.3%) | N/A |
| — Left-sided colitis (E2) | 11 (68.8%) | N/A |
| — Extensive colitis (E3) | 4 (25.0%) | N/A |
| Biologic-naïve | 12 (75.0%) | N/A |
| Endoscopically active disease | 16 (100%) | N/A |
| SCCAI | 6.5 (3.5-8.3) | N/A |

